# A CRISPR/Cas9 based strategy to manipulate the Alzheimer’s amyloid pathway

**DOI:** 10.1101/310193

**Authors:** Jichao Sun, Jared Carlson-Stevermer, Utpal Das, Minjie Shen, Marion Delenclos, Amanda M. Snead, Lina Wang, Jonathan Loi, Andrew J Petersen, Michael Stockton, Anita Bhattacharyya, Mathew V. Jones, Andrew A. Sproul, Pamela J. McLean, Xinyu Zhao, Krishanu Saha, Subhojit Roy

## Abstract

The gradual accumulation of amyloid-β (Aβ) is a neuropathologic hallmark of Alzheimer’s disease (AD); playing a key role in disease progression. Aβ is generated by the sequential cleavage of amyloid precursor protein (APP) by β- and γ-secretases, with BACE-1 (β-site APP cleaving enzyme-1) cleavage as the rate limiting step ^1–3^. CRISPR/Cas9 guided gene-editing is emerging as a promising tool to edit pathogenic mutations and hinder disease progression ^4,5,6^ However, few studies have applied this technology to neurologic diseases ^7–9^. Besides technical caveats such as low editing efficiency in brains and limited in vivo validation ^7^, the canonical approach of ‘mutation-correction’ would only be applicable to the small fraction of neurodegenerative cases that are inherited (i.e. < 10% of AD, Parkinson’s, ALS); with a new strategy needed for every gene. Moreover, feasibility of CRISPR/Cas9 as a therapeutic possibility in sporadic AD has not been explored. Here we introduce a strategy to edit endogenous APP at the extreme C-terminus and reciprocally manipulate the amyloid pathway – attenuating β-cleavage and Aβ, while up-regulating neuroprotective a-cleavage. APP N-terminus, as well as compensatory APP homologues remain intact, and key physiologic parameters remain unaffected. Robust APP-editing is seen in cell lines, cultured neurons, human embryonic stem cells/iPSC-neurons, and mouse brains. Our strategy works by limiting the physical association of APP and BACE-1, and we also delineate the mechanism that abrogates APP/BACE-1 interaction in this setting. Our work offers an innovative ‘cut and silence’ gene-editing strategy that could be a new therapeutic paradigm for AD.

---

Our broad idea is to rationally edit small segments of wild-type (WT) proteins known to play key roles in the progression of sporadic disease, with the ultimate goal of attenuating their pathologic activity. As endogenous proteins expectedly play physiologic roles as well, it is also important to conserve the normal function of these molecules, as far as possible. Motivated by this idea, we designed sgRNAs targeting the extreme C-terminus of mouse APP, and one of these led to robust APP-editing, as shown in **Fig. 1** (see protospacer adjacent motif – PAM – site and genomic target recognized by the sgRNA in **Fig. 1a**). APP-editing resulted in attenuated staining with an antibody (Y188) recognizing the extreme C-terminus of APP that is distal to the sgRNA-targeting site (neuroblastoma cells shown in **Fig. 1b**, see Y188 epitope in **Fig. 1a**). APP-sgRNA also attenuated the production of APP C-terminal fragments (CTFs; **Fig. 1c, d** - time-course of editing in **Fig. 1e**). However, an antibody recognizing the APP N-terminus (22C11) showed no differences between control and sgRNA-treated samples (**Fig. 1c**), suggesting that the editing only affected the short intracellular C-terminus. Genomic deep-sequencing confirmed efficient editing of mouse APP at the expected target (**Fig. 1f**).

**Fig. 1.**
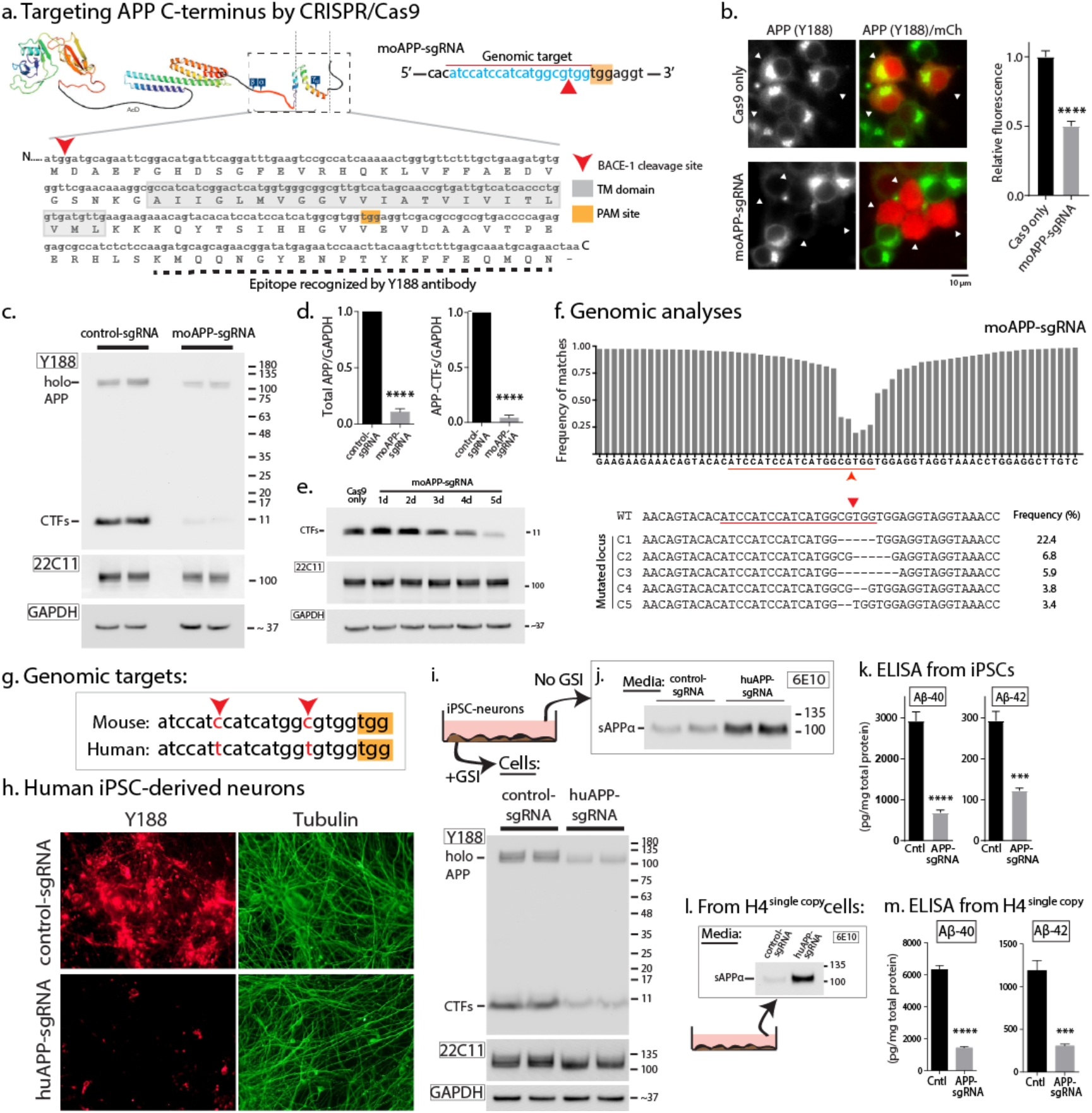
Strategy and manipulation of the amyloid pathway by CRISPR/Cas9 editing. **(a)** Schematic and C-terminal sequence of mouse APP showing PAM site and sgRNA target. Note that the Y188 antibody recognizes the last 20 aa [schematic adapted from (*16*), permission pending]. **(b)** Neuro2A cells were transfected with APP-sgRNA and Cas9 (or Cas9 only), and immunostained with the Y188 antibody (after 5 days in culture). The mCherry protein was co-expressed with CRISPR/Cas9 to label transfection positive cells. Note decreased APP fluorescence, indicating APP editing (quantified on right, mean ± SEM of ~40 cells from two independent experiments). **(c and d)** Neuro2A cells were transduced by lentiviral vectors carrying APP-sgRNA and Cas9 (or non-targeting control-sgRNA/Cas9 as control) and immunoblotted with Y188 and 22C11 antibodies (latter recognizes APP N-terminus). A gamma secretase inhibitor (GSI) was added to allow detection of accumulated APP CTF’s (see methods, GAPDH used as loading controls). Note attenuation of APP-CTFs in APP-sgRNA treated samples, but no change in 22C11 signal. Blots quantified in (d), mean ± SEM of three independent experiments. **(e)** Time course of APP-editing in neuro2a cells. Cells were transfected with a vector carrying APP-sgRNA and Cas9, and APP-CTFs were analyzed by Western blotting (in the presence of GSI). **(f)** Deep sequencing of APP C-terminus in neuro2A cells. Top: Frequency of base-pair matches between gRNA-edited and WT mouse sequence. Red underline marks the sgRNA target sequence and arrowhead denotes predicted cut-site. Note extensive mismatch around predicted cut-site, indicating robust editing. Bottom: Major mutated APP loci resulting from sgRNA-editing, and their frequencies. **(g)** Comparison of mouse and human APP-sgRNA targeting sequences (red arrowheads indicate nucleotide differences; yellow bar denotes the PAM site). **(h)** Human iPSC derived NPCs were transduced by lentiviral vectors carrying APP-sgRNA and Cas9 (or non-targeting control-sgRNA/Cas9 as control) and differentiated into neurons. After 3 weeks for differentiation, cells were immunostained with the Y188 and Tuj1 antibodies. Note decreased APP fluorescence, indicating APP editing. **(i)** Samples from iPSC derived neurons above (transduced with control-sgRNA/Cas9 or APP-sgRNA/Cas9) were immunoblotted with the Y188 and 22C11 antibodies (in the presence of GSI). Note attenuation of APP blotting with Y188 in cells carrying the APP-sgRNA. **(j)** Media from iPSC derived neurons above were immunoblotted for extracellular sAPPα with 6E10 antibody (in the absence of GSI). Note increase in APP *α*-cleavage in the APP-sgRNA treated sample. **(k)** ELISA of media from iPSC derived neurons (treated as above). Note decreased Aβ in the APP-sgRNA treated samples (mean ± SEM of three independent experiments). **(l)** The single copies of APP and BACE-1 were integrated into H4 genome, and the generated H4^single copy^ cell line was transduced with lentiviral vectors carrying control-sgRNA/Cas9 or APP-sgRNA/Cas9 (see Extended Data Figure 2 and methods). Media were immunoblotted for extracellular sAPP*α* (6E10 antibody). Note increase in APP *α*-cleavage in the APP-sgRNA treated sample. **(m)** ELISA of media from the H4^single copy^ cell line (treated as above). Note decreased Aβ in the APP-sgRNA treated samples (mean ± SEM of three independent experiments).

Though the abovementioned TGG PAM is conserved in both mouse and human APP, the upstream targeting sequence differs only by two nucleotides (**Fig. 1g**). Despite this, the mouse APP-sgRNA was unable to edit human APP, perhaps reflecting the specificity of the CRISPR/Cas9 system; also attested by other groups ^10–12^. However, a sgRNA specific to the human APP targeting sequence robustly edited APP in HEK293 (**Extended Data Fig. 1a-c**), as well as in human embryonic stem cells (**Extended Data Fig. 1d-f**); as determined by immunostaining, western blots and deep-sequencing. APP editing and decreased CTFs was also seen in human iPSCs differentiated to neurons (**Fig. 1h, i**). While APP cleavage by BACE-1 initiates the cascade of events leading to Aβ deposition, the alternative a-cleavage “anti-amyloidogenic” pathway – mediated by a-secretases – is thought to be protective [reviewed in *13*]. Interestingly, sAPPa was up-regulated in iPSC-neurons (**Fig. 1j**), suggesting activation of the neuroprotective a-cleavage pathway. ELISAs showed corresponding attenuation of Aβ-40/42 in the iPSC-neurons, confirming inhibition of β-cleavage (**Fig. 1k**).

The above data suggest that the APP-sgRNA has reciprocal effects on APP a- and β-cleavage. To determine effects of the sgRNA in a more controlled setting, we engineered a stable H4 neuroglioma cell line expressing single copies of APP and BACE-1, using the FlpIn system (APP/BACE^single_copy^; for details, see **Extended Data Fig. 2a** and methods). Endogenous APP/BACE-1 levels are negligible in these cells, thus almost all the APP/BACE-1 expression is from the introduced single-copies (see **Extended Data Fig. 2b**).

We also tagged APP and BACE-1 in these cells to the N- and C-terminal fragments of the Venus fluorescent protein (VN/VC) respectively – as in our previous study ^14^ – for two reasons. First, fluorescence complementation of APP:VN and BACE-1:VC reflects the physical association of this substrate-enzyme pair ^14^, allowing us to directly evaluate editing efficiency by monitoring YFP fluorescence. Second, the APP-CTFs generated from APP:VN are easier to identify in Western blots (as they run higher). Indeed, transduction of APP/BACE^single_copy^ cells with a lentivirus carrying APP-sgRNA and Cas9 essentially eliminated Venus complementation (**Extended Data Fig. 2c, d**), confirming robust APP editing. Biochemical analyses using APP antibodies against extra- or intra-cellular epitopes confirmed that the sgRNA specifically inhibited APP β-cleavage (**Extended Data Fig. 2e, f**). Indeed, in line with the data from iPSC-neurons, the sgRNA had reciprocal effects on sAPPα and Aβ **(Fig. 11, m);** providing confidence that our gene editing strategy is favorably manipulating the amyloid pathway.

Off-target effects of CRISPR/Cas9 is a potential concern. Towards this, we asked if our mouse and human APP-sgRNA were able to edit the top five computationally predicted off-target sites **(Extended Data Fig. 3a;** also see **Extended Data Table 1**). No editing was detected **(Extended Data Fig. 3b-e),** further attesting specificity. Though APP null mice are viable with minor deficits, there is compensation by the two APP homologues APLP1 and 2 that undergo similar processing as APP (reviewed in *15,16*). APLP1 and 2 were not amongst the top 50 predicted off-target sites, as their corresponding sgRNA-target sites were substantially different from APP (see sequences in **Extended Data Fig. 3f**). For further assurance that our sgRNA was not editing the APP homologues, we performed TIDE off-target analyses ^*17*^ on cells carrying the sgRNA. As shown in **Extended Data Fig. 3g**, TIDE analyses showed no editing of APLP 1/2 by the sgRNA.

APP has physiologic roles in axon growth and signaling [reviewed in *18*]. As noted above, the N-terminus of APP – thought to play roles in axon growth and differentiation – is entirely preserved in our setting. However, the C-terminal APP intracellular domain (AICD) has been implicated in gene transcription, though the effect appears to be both physiologic and pathologic ^19,20^. To examine potential deleterious effects of editing the extreme C-terminus of APP, we turned to cultured hippocampal neurons where various parameters like neurite outgrowth and synaptic structure/function can be confidently evaluated. For these studies, we generated AAV9 viruses carrying the APP-sgRNA and Cas9, tagged with GFP and HA respectively (see vector design in **Fig. 2a**) that transduced almost all cultured neurons (**Fig. 2b** and data not shown). In blinded analyses, we found no significant effect of the APP-sgRNA on neurite outgrowth, axon-length, synaptic organization, or neuronal activity (**Fig. 2**). Although further studies are needed, we note that besides editing only a small segment of APP, our strategy: 1) does not completely block β-cleavage; and 2) does not affect the physiologic processing of APLP 1/2 (that also generate CTFs).

**Fig. 2.**
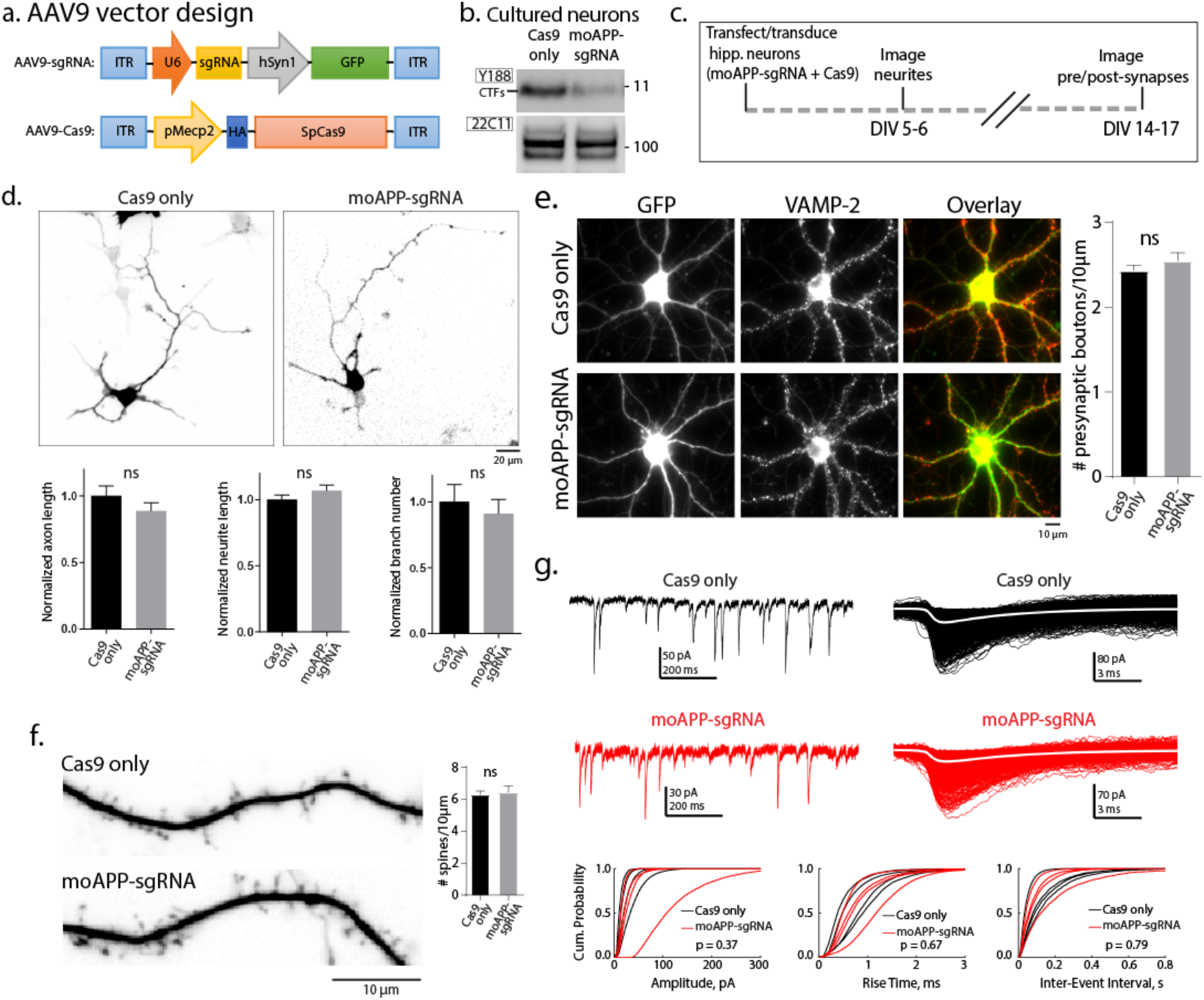
Effect of APP C-terminus editing on neuronal physiology. **(a)** AAV9-sgRNA and AAV9-Cas9 expression vectors. Note that the sgRNA vector co-expresses GFP and the Cas9 is tagged to HA, for identification of transduced neurons. **(b)** Cultured hippocampal neurons were transduced with AAV9s carrying APP-sgRNA/Cas9 (or Cas9 only) and immunoblotted with the Y188 and 22C11 antibodies (in the presence of GSI). Note attenuation of CTFs by the APP-sgRNA. **(c)** Neurons were transfected (at the time of plating) with a vector expressing APP-sgRNA and Cas9. Neuritic/axon outgrowth was analyzed after 5-6 days, and pre/post-synapse was analyzed after 14-17 days. **(d)** Top: Representative images of neurons transfected with the APP-sgRNA/Cas9 (or Cas9 alone). Bottom: Axon length and number of neurites/branches in the APP-sgRNA/Cas9 (or Cas9 alone) groups; note that there was no significant difference (mean ± SEM of ~30 cells from two independent experiments). **(e)** Neurons were infected with AAV9 viruses carrying APP-sgRNA/Cas9 (or Cas9 only as controls), and fixed/stained with the presynaptic marker VAMP2. Note that the presynaptic density (VAMP2 puncta) was similar in both groups (quantified on right, mean ± SEM of VAMP2 staining along ~ 25 dendrites from two independent experiments). **(f)** Neurons were transfected with APP-sgRNA/Cas9 (or Cas9 only as controls). Spine density in the APP-sgRNA/Cas9 (or Cas9 only) groups was also similar, quantified on right (mean ± SEM of ~16 dendrites from two independent experiments). **(g)** Miniature excitatory postsynaptic currents (mEPSC) were recorded from neurons infected with AAV9-APP-sgRNA/Cas9 or AAV9-Cas9 alone. Top: Representative mEPSC traces in control and APP-sgRNA transduced neurons. Corresponding alignments of mEPSCs with average (white traces) are shown on right. Bottom: Cumulative histograms of mEPSC amplitude, 20-80% rise-time and inter-event interval in APP-sgRNA/Cas9 and the Cas9-only infected neurons (note no significant differences).

Next we asked if our sgRNA could edit APP in vivo. Injection of the AAV9s into mouse hippocampi (**Fig. 3a**) led to efficient transduction of both sgRNA and Cas9 in dentate neurons (86.87 ± 2.83 % neurons carrying the sgRNA also had Cas9 – sampling of 495 neurons from 3 brains; see representative images in **Fig. 3b**). Immunostaining of transduced neurons with the APP Y188 antibody suggests editing of endogenous APP in vivo (**Fig. 3c**). To achieve a more widespread expression of the sgRNA and Cas9 in mouse brains – and also evaluate editing by biochemistry – we injected the viruses into the ventricles of neonatal (P0) mice and examined the brain after 2-4 weeks (**Fig. 3e**). Previous studies have shown that when AAVs are injected into the ventricles of neonatal mice, there is widespread delivery of transgenes into the brain – also called somatic transgenesis ^21,22^. Indeed, we saw patchy attenuation of APP Y188 staining in cortical regions (**Fig. 3f**), and also a decrease in CTFs by western blotting (**Fig. 3g, h**); suggesting that our gRNA can edit APP in vivo.

**Fig. 3.**
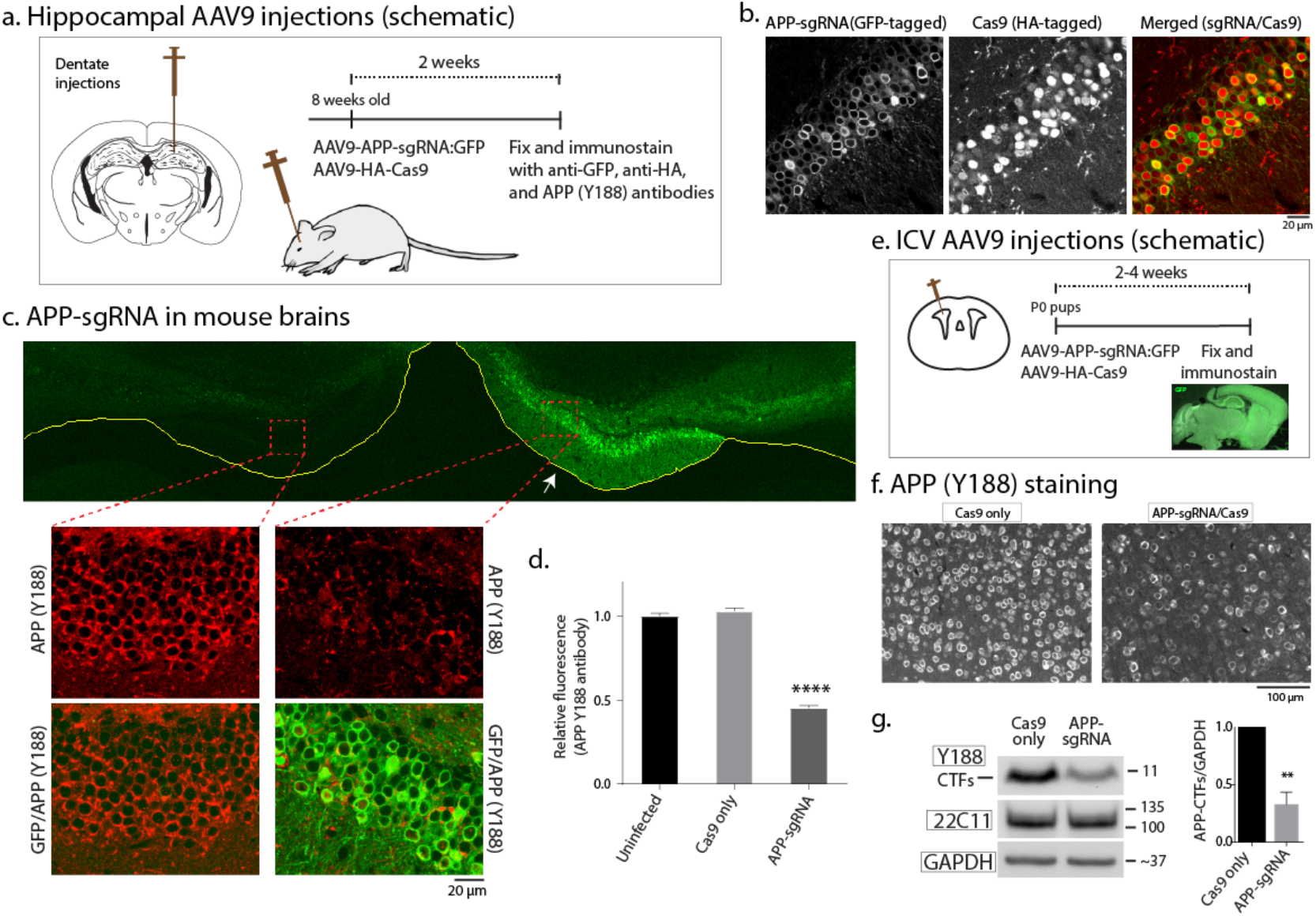
Editing of APP C-terminus in vivo. **(a)** AAV9-sgRNA and AAV9-Cas9 were stereotactically co-injected into dentate gyms of 8-week old mouse brains (bottom). Two weeks after viral delivery, brains were perfused, fixed, and immunostained with anti-GFP, anti-HA and anti-APP(Y188) antibodies, **(b)** Co-expression of AAV9-sgRNA-GFP and AAV9-HA-Cas9 in the dentate gyrus. Note that majority of neurons are positive for both GFP and HA (~ 87 % of the cells were positive for both; sampling from 3 brains), **(c and d)** Coronal section of a mouse hippocampi injected on one side (marked by arrow) with the AAV viruses as described above. Note attenuated Y88 staining of neurons on the injected side, indicating APP-editing. Fluorescence quantified in (d), mean ± SEM, data from three brains, **(e)** Intracerebroventrical injection of the AAV9 viruses into PO pups. Note widespread delivery of gRNA into brain, as evident by GFP fluorescence, **(f)** Brain sections from above were immunostained with the Y188 antibody. Note attenuated Y188 staining in the APP-sgRNA/Cas9 transduced sample, suggesting APP-editing. **(g)** Western blots of the brains from (e). Note decreased expression of CTFs in the APP-sgRNA/Cas9 transduced brains; blots quantified on right.

Finally, we sought to understand the mechanism by which CRISPR-mediated APP editing attenuates the β-cleavage pathway (note that the sgRNA-editing site is distant from the β-cleavage site). Genomic sequencing showed that sgRNA-editing leads to a translational product where the last 36 aa of APP are truncated (**Fig. 4a**). To evaluate functional consequences of editing, we generated a truncated “APP CRISPR-mimic” construct (APP-659). Using our fluorescence complementation assay ^14^, we first asked if APP-659 interacted with BACE-1. APP-659/BACE-1 approximation was greatly attenuated (**Fig. 4b**), along with a decrease in β-CTF generation (**Fig. 4c**). What is the mechanism underlying this decrease in APP/BACE-1 complementation? Since the sgRNA targeting site is distant from the BACE-1-cleavage site (see **Fig. 1a**), it seems unlikely that gene editing directly interferes with APP/BACE-1 interaction. Instead, trafficking alterations of the edited APP molecules – eventually leading to decreased APP/BACE-1 approximation – seem more likely. APP is synthesized in the ER→Golgi pathway, and Golgi-derived vesicles carrying APP are transported into axons and dendrites, inserting into the plasma membrane. Subsequently, surface APP is internalized into endosomes, where it is cleaved by BACE-1, and this is thought to initiate β-cleavage ^23–5^.

**Fig. 4.**
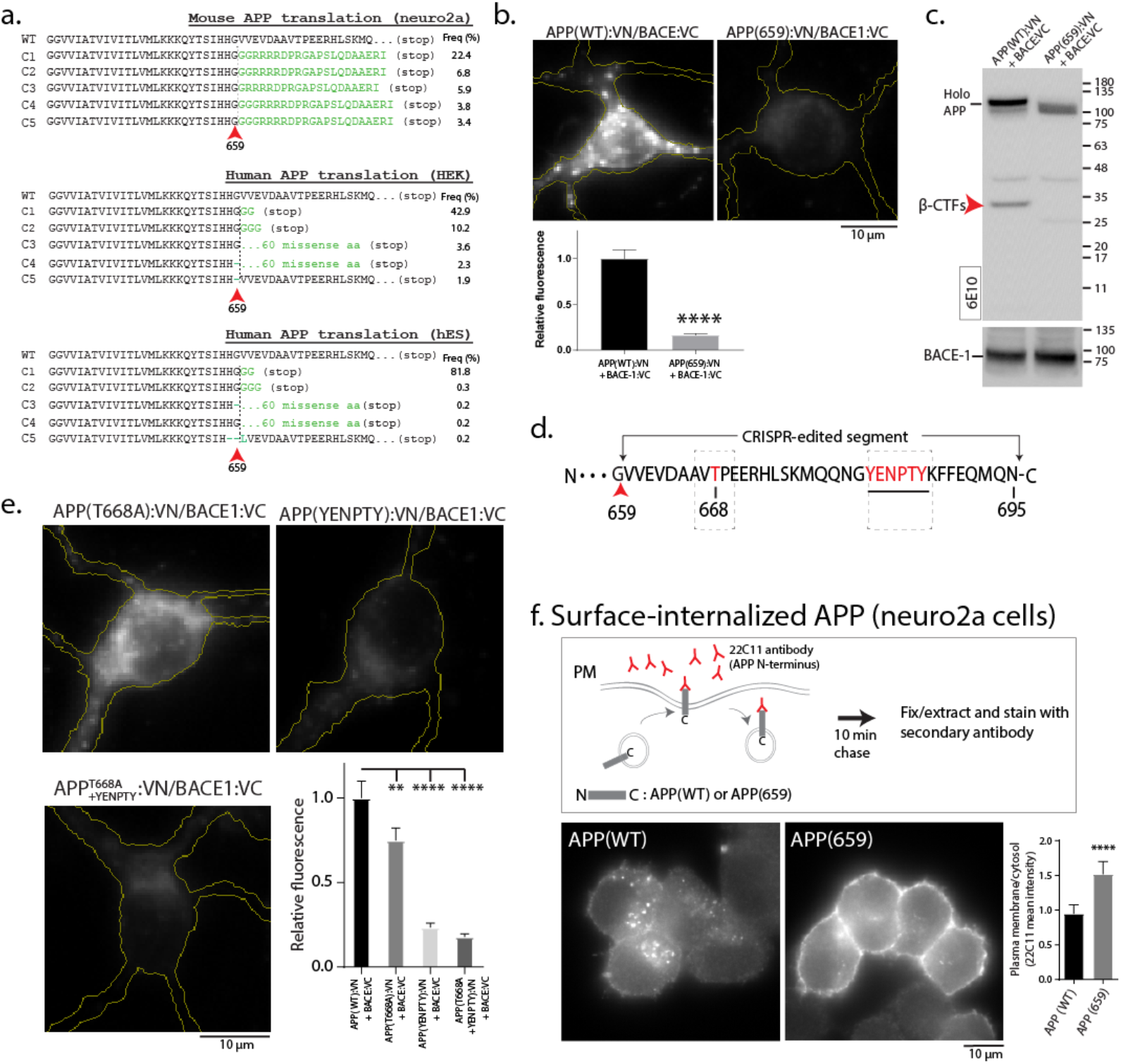
Mechanism of CRISPR-guided APP editing. **(a)** Predicted APP translational products after CRISPR/Cas9 editing in mouse (neuro2A) and human (HEK and human ES) cells for the major mutant alleles observed in deep sequencing. Note that after editing, APP is translated up-to amino acid 659 (red arrowheads). **(b)** APP/BACE-1 interaction – as evaluated by fluorescence complementation in cultured hippocampal neurons – was attenuated in neurons transfected with a “CRISPR-mimic” APP (APP659:VN; quantified below, mean ± SEM of ~35 cells from three independent experiments). **(c)** APP β-cleavage is attenuated in cells transfected with APP659. HEK cells were co-transfected with APPWT (or APP659) tagged to VN, and BACE-1:VC; and immunoblotted with the 6E10 antibody. Note decreased β-CTFs in cells carrying the truncated APP plasmid. **(d)** Schematic showing the CRISPR-edited C-terminus portion of APP. Note that the threonine at 668 position, and the endocytic YENPTY motif (dashed boxes) are thought to play roles in Aβ production. **(e)** APP/BACE-1 interaction – as evaluated by fluorescence complementation in cultured hippocampal neurons – was most markedly attenuated in neurons transfected with mutant YENPTY. **(f)** Strategy of APP internalization assay. Neuro 2a cells are transfected with APP:GFP or APP659:GFP. After incubation with anti N-terminal APP antibody (22C11) for 10 min, the cells were fixed and stained with secondary antibody to visualize the cell surface and internalized APP. Note the cell surface accumulation and decreased internalization of the CRISPR-mimic APP659.

Accordingly, we systematically explored various trafficking steps in hippocampal neurons. First, we visualized axonal and dendritic transport of APP-WT and APP-659. Although there were modest changes (**Extended Data Fig. 4** and **Extended Data Table 2**), it seems unlikely that such transport perturbations would lead to the dramatic attenuation of β-cleavage and Aβ-production seen in our experiments. Asking if amino-acid residues within the CRISPR-edited segment of APP might offer clues, we noted that APP residues T668 and Y682-Y687 (the “YENPTY motif”, see **Fig. 4d**; also present in APLP1/2) in the edited segment have been reported to play a role in Aβ production ¾^27^. Specifically, APP phosphorylated at T668 preferentially colocalizes with BACE-1 in endosomes ^26^, and the YENPTY motif mediates APP internalization from the plasma membrane ^28^. Indeed, the extent of APP/BACE-1 attenuation by the YENPTY mutation strongly resembled the decrease in fluorescence complementation by the APP-659 (“CRISPR-mimic”) construct (**Fig. 4e**). A prediction from these experiments is that the endocytosis of APP-659 from the cell surface should be attenuated; and indeed, this was the case in neuronal internalization assays (**Fig. 4f**). Collectively, the data suggest that our gene-editing approach does not have a major effect on post-Golgi trafficking of APP, but attenuates its endocytosis from cell surface, and consequently, its interaction with BACE-1 in endosomes. Since most of the APP α-cleavage is thought to occur at the cell surface ^29^, this may also explain why the non-amyloidogenic pathway is enhanced by our approach.

Using CRISPR/Cas9 technology, here we provide proof of concept for a gene-editing strategy that can favorably manipulate the amyloid pathway – attenuating β-cleavage and Aβ production, while up-regulating the neuroprotective α-cleavage. APP editing was efficient in a variety of human and mouse cells, neurons, and in vivo. After almost thirty years of controversies and failures, gene therapy of neurologic diseases has turned a page, with striking results in clinical trials ^6,30^. A key advance has been the development of AAV9-based vectors that can diffusely deliver genes throughout the nervous system ^31^, and our vision is to leverage these innovations to develop gene-editing therapies for AD. In principle, our ‘cut and silence’ CRISPR-editing approach might also work for attenuating other endogenous pathology-driving proteins in neurodegenerative diseases – such as BACE, tau, presenilins, α-synuclein, and TDP-43 – and may usher in a new way of therapeutically tackling these devastating diseases.

## EXPERIMENTAL PROCEDURES

### Constructs, antibodies and reagents

For transient co-expression of CRISPR/Cas9 components, APP gRNA nucleotides were synthesized and cloned into pU6-(Bbs1)_CBh-Cas9-T2A-mCherry vector at Bbs1 site. For viral transduction, a dual vector system was used to deliver CRISPR/Cas9 components using AAV9 ^32^. For making the AAV9 vectors, the APP gRNA was cloned into pAAV9-U6sgRNA(SapI)_hSyn-GFP-KASH-bGH vector at Sap1 site. The CRISPR/Cas9 stable cell lines were generated by lentivirus infection as follows. The APP gRNA was cloned into lentiCRISPR v2 vector at Bbs1 site to produce lentivirus ^33^. For making APP deletions and relevant constructs, the human APP659 truncation was PCR amplified and cloned at Hind3 and Sac2 sites of pVN to generate pAPP659:VN. The BBS-APP659 was PCR amplified and cloned into pBBS-APP:GFP at Hind3 and Sac2, replacing BBS-APP, to generate pBBS-APP659:GFP. The pBBS-APP^YENPTY^:GFP was generated by site directed mutagenesis from pBBS-APP:GFP. The pAPP^T668A^:VN and pAPP^T668A+YENPTY^:VN were generated by site directed mutagenesis from pAPP:VN and pAPP^YENPTY^:VN. Antibodies used were as follows: APP Y188 (ab32136; Abcam), APP 22C11 (MAB348; Millipore), APP 6E10 (803001; BioLegend), BACE-1 (MAB931; R&D), GAPDH (MA5-15738, ThermoFisher), GFP (ab290, Abcam), GFP (A10262, Invitrogen), HA (901513, BioLegend), VAMP2 (104211, Synaptic Systems). Reagents were as follows: a-bungarotoxin Alexa-594 conjugate (Life Technologies), Tubocurarine chloride (Sigma), γ-secretase inhibitor BMS-299897 (Sigma), Rho Kinase (ROCK)-inhibitor H-1152P (Calbiochem) and Dynasore (Sigma).

### Cell cultures, transfections, viral production/infections, and biochemistry

HEK293 and neuro2a cells (ATCC) were maintained in DMEM with 10% FBS. Cells were transfected with Lipofectamine 2000 and collected 5 days after transfection for biochemical and immunostaining analysis. Primary hippocampal neurons were obtained from postnatal (P0-P1) CD1 mice (either sex), and transiently transfected using Lipofectamine 2000 or Amaxa 4D system (Lonza). Dissociated neurons were plated at a density of 30,000 cells/cm^2^ on poly-D-lysine-coated glass-bottom culture dishes (Mattek) and maintained in Neurobasal/B27 medium with 5% CO2. For APP/BACE-1 interaction, APP internalization and APP transport studies, DIV 7 neurons were cultured for ~18-20 h after transfection. For spine density analysis, DIV7 neurons were transfected with soluble markers and cultured for 7 d before imaging. For testing the effect of CRISPR/Cas9 on neuronal development, neurons were electroporated with the respective constructs before plating using an Amaxa 4D-Nucleofector system with the P3 Primary Cell 4D-Nucleofector X kit S and program CL-133.

For western blotting and electrophysiology, DIV7 cultured neurons were infected with either AAV9-APP gRNA-GFP (2.24x10^13^ Vg/ml) and AAV9-Cas9 (2.4x10^14^ Vg/ml), or AAV9-GFP (2.58x10^13^ Vg/ml) and AAV9-Cas9 at a multiplicity of infection (MOI) of 1.5x10^5^. Neurons were analyzed 7 days post-infection. Lentivirus was produced from HEK293FT cells as described ^34^. Briefly, HEK293FT cells (Life Technologies) were maintained in DMEM with 10% FBS, 0.1mM NEAA, 1 mM sodium pyruvate and 2mM Glutamine. Cells were transfected with lentiviral-target and helper plasmids at 80-90% confluency. 2 days after transfection, the supernatant was collected and filtered with 0.45 μm filter. For experiments with hESCs, cells were cultured on a Matrigel substrate (BD Biosciences) and fed daily with TeSR-E8 culture media (StemCell Technologies). When the cells were around 60-70% confluent, they were infected with a 50/50 mixture of TeSR-E8 (with 1.0 μM H-1152P) and lentivirus supernatant. After 24 h, the virus was removed, and the cells were fed for 2 days (to recover). After 3 days, cells were treated with 0.33 μg/mL of puromycin for 72 h to select for virally-integrated hESCs. For HEK and neuro2a cell lines, cells were infected with the lentivirus carrying APP-sgRNA and Cas9 for 24 h. And then cells were fed for 1 day to recover. After 2 days, cells were treated with 1 μg/mL of puromycin for 72 h to select for virally-integrated cells.

Human NPCs were generated as has been described previously ^35^, using manual rosette selection and Matrigel (Corning) to maintain them. Concentrated lentiviruses express control-sgRNA or APP-sgRNA were made as described previously ^36^, using Lenti-X concentrator (Clontech). The NPCs were transduced with either control-sgRNA or APP-sgRNA after Accutase splitting, and were submitted to puromycin selection the subsequent day. Polyclonal lines were expanded, and treated with puromycin for 5 more days before banking. Neuronal differentiations were carried out by plating 165,000 cells/12 well-well in N2/B27 media (DMEM/F12 base) supplemented with BDNF (20 ng/mL; R&D) and laminin (1 ug/mL; Trevigen).

For biochemistry, cell lysates were prepared in PBS + 0.15% Triton X-100 or RIPA supplemented with protease inhibitor cocktail, pH 7.4. After centrifuging at 12,000 g for 15 min at 4 °C, supernatants were quantified and resolved by SDS-PAGE for western blot analysis. For sAPPa detection, cell culture medium was collected and centrifugated at 2,000 g for 15 min at RT. The supernatants were resolved by SDS-PAGE for western blot analysis; band intensities were measured by ImageJ. Human Aβ40 and Aβ42 were detected using kits, according to the manufacturer’s instructions (Thermo KHB3481 and KHB3544). Briefly, supernatants from H4^single copy^ cells or human iPSC derived neurons were collected and diluted (x5 for H4 and x2 for iPSC-neuron). The diluted supernatants and the human Aβ40/42 detection antibodies were then added into well, and incubated for 3 h at RT with shaking. After washing (x4), the anti-Rabbit IgG HRP solution was added and incubated for 30 min at RT. The stabilized Chromogen was added after washing (x4), and incubated for another 30 min at RT in the dark. After addition of stop solution, absorbance at 450 nm was read using a luminescence microplate reader.

### Developing a single-copy, stable APP/BACE-1 cell line

H4 tetOff FlpIn empty clone was maintained in OptiMEM with 10% FBS, 200 μg/mL G418 and 300 μg/mL Zeocin. To generate an APP:VN/BACE-1:VC stable cell line carrying single copies of APP and BACE-1, the expressing plasmid and pOG44 plasmids were transfected with Lipofectamine 2000. 2 days after transfection, cells were selected with 200 μg/mL hygromycin B and 200 μg/mL G418 for 1 week. A monoclonal cell line with stable expression was selected by cell sorting, based on fluorescence-complementation of the tagged VN/VC fragments. H4 stable cell lines were then infected with the lentivirus carrying APP-gRNA and Cas9, as described above. After 24 h, the virus was removed, and cells were fed for 1 day to recover. After 2 days, cells were treated with 0.7 μg/mL of puromycin for 72 h to select for virally-integrated cells.

### Immunofluorescence, microscopy/image analysis, APP trafficking and endocytosis assays

For immunostaining of endogenous APP or VAMP2, cells were fixed in 4% PFA/sucrose solution in PBS for 10 min at room temperature (RT), extracted in PBS containing 0.2% Triton X-100 for 10 min at RT, blocked for 2 h at RT in 1% bovine serum albumin and 5% FBS, and then incubated with rabbit anti-APP (1:200) or mouse anti-VAMP2 (1:1000) diluted in blocking buffer for 2 h at RT. After removal of primary antibody, cells were blocked for 30 min at RT, incubated with goat anti–rabbit (Alexa Fluor 488) or goat anti–mouse (Alexa Fluor 594) secondary antibody at 1:1000 dilution for 1 h at RT and then mounted for imaging. z-stack images (0.339 μm z-step) were acquired using an inverted epifluorescence microscope (Eclipse Ti-E) equipped with CFI S Fluor VC 40× NA 1.30 (Nikon). An electron-multiplying charge-coupled device camera (QuantEM: 512SC; Photometrics) and LED illuminator (SPECTRA X; Lumencor) were used for all image acquisition. The system was controlled by Elements software (NIS Elements Advanced Research). z-stacks were subjected to a maximum intensity projection. For APP Y188 staining, the average intensity of single cell body (neuro2A, HEK293 and neurons) or the whole colony (hESCs) was quantified. All the images were analyzed in Metamorph and ImageJ.

Spine density experiments were done as described previously ^37^. Briefly, DIV 7 neurons were transfected with desired constructs for 7 days, and secondary dendrites were selected for imaging. z-stack images were captured using a 100x objective (0.2 μm z-step) and subjected to a maximum intensity projection for analysis. For the APP/BACE-1 complementation assay, DIV 7 neurons were transfected with desired constructs for ~15-18 h and fixed. z-stack images were captured using a 40x objective (0.339 μm z-step) and subjected to a maximum intensity projection. The average intensity within cell bodies was quantified.

For trafficking studies in axons and dendrites, imaging parameters were set at 1 frame/s and total 200 frames. Kymographs were generated in MetaMorph, and segmental tracks were traced on the kymographs using a line tool. The resultant velocity (distance/time) and run length data were obtained for each track, frequencies of particle movements were calculated by dividing the number of individual particles moving in a given direction by the total number of analyzed particles in the kymograph, and numbers of particles per minute were calculated by dividing the number of particles moving in a given direction by the total imaging time.

APP endocytosis assay was done as described previously ^38^. Cells expressing APP-GFP or APP659-GFP were starved with serum-free medium for 30 min and incubated with anti-APP (22C11) in complete medium with 10 mM HEPES for 10 min. And then, cells were fixed, permeablized and immunostained for 22C11. The mean intensity of 22C11 along plasma membrane was calculated by dividing the total intensity along plasma membrane (= intensity of whole cell – intensity of cytoplasm) with area of plasma membrane (= area of whole cell – area of cytoplasm). The ratio of mean intensities between plasma membrane and cytoplasm was quantified.

### Stereotactic injection of AAV9s into the mouse brain and histology

In vivo injection and immunofluorescence staining was done as described previously ^39^. Briefly, 1.5๑1 of 1:2 AAV9 mixture of AAV9-APP gRNA-GFP (or AAV9-GFP) and AAV9-Cas9 was injected into the dentate gyrus (-2.0, ±1.6, -1.9) of 8-week old male C57BL/6 mice (either sex). 2-weeks after surgery, the mice were sacrificed by trans-cardiac perfusion of saline, followed by 4% PFA. The brains were dissected, post-fixed with 4% PFA overnight, immersed in 30% sucrose until saturation, and sectioned at 40 μm. Sections were immunostained with following antibodies: mouse anti-HA (1:1000, BioLegend, clone 16B12), chicken anti-GFP (1:1000, Invitrogen, polyclonal) and rabbit anti-APP (1:200, Abcam, clone Y188). Images were acquired using Zeiss LSM800 confocal microscope. Average intensities of APP staining in cell bodies was quantified using Metamorph.

### Intracerebroventricular injections and histology

All animal procedures were approved by the Mayo Institutional Animal Care and Use Committee and are in accordance with the NIH Guide for Care and Use of Laboratory animals. Free hand bilateral intracerebroventricular (ICV) injections were performed as previously described ^40^ in C57BL/6J mouse pups. On post-natal day 0, newborn pups were briefly cryoanesthetized on ice until no movement was observed. A 30-gauge needle attached to a 10 μl syringe (Hamilton) was used to pierce the skull of the pups just posterior to bregma and 2 mm lateral to the midline. The needle was held at a depth of approximately 2 millimeters, and 2 μl of a mixture of AAV9 viruses (ratio 1:2 of AAV9--APP gRNA-GFP or AAV9-GFP+ AAV9-Cas9) were injected into each cerebral ventricle. After 5 minutes of recovery on a heat pad, the pups were returned into their home cages. Mice were sacrificed 15 days after viral injection. Animals were deeply anesthetized with sodium pentobarbital prior to transcardial perfusion with phosphate buffered saline (PBS), and the brain was removed and bisected along the midline. The left hemisphere was drop-fixed in 10% neutral buffered formalin (Fisher Scientific, Waltham, MA) overnight at 4°C for histology, whereas the right hemisphere of each brain was snap-frozen and homogenized for biochemical analysis. Formalin fixed brains were embedded in paraffin wax, sectioned in a sagittal plane at 5-micron thickness, and mounted on glass slides. Tissue sections were then deparaffinized in xylene and rehydrated. Antigen retrieval was performed by steaming in distilled water for 30 min, followed by permeabilization with 0.5% Triton-X, and blocking with 5% goat serum for 1 hour. Sagittal sections were then incubated with primary anti-GFP antibody (1:250, Aves, chicken polyclonal) and anti-APP antibody (1:200, Abcam, clone Y188) overnight at 4°C. Sections were incubated with the secondary antibodies Alexa 488-goat anti-chicken and Alexa 568-goat anti rabbit (1:500, Invitrogen) for 2h at room temperature. Sections were washed and briefly dipped into 0.3% Sudan Black in 70% ethanol prior to mounting.

### Electrophysiology

A coverslip with cultured cells at a density of 60,000 cells/cm^2^ was placed in a continuously perfused bath, viewed under IR-DIC optics and whole-cell voltage clamp recordings were performed (-70 mV, room temp.). The extracellular solution consisted of (in mM): 145 NaCl, 2.5 KCl, 1 MgCl2, 2 CaCl2, 10 HEPES and 10 dextrose, adjusted to 7.3 pH with NaOH and 320 mOsm with sucrose. Whole-cell recordings were made with pipette solutions consisting of (in mM) 140 KCl, 10 EGTA, 10 HEPES, 2 Mg2ATP and 20 phosphocreatine, adjusted to pH 7.3 with KOH and 315 mOsm with sucrose. Excitatory synaptic events were isolated by adding 10 μM bicuculline to block GABA (subscript A) receptors. Miniature synaptic events were isolated by adding 100 nM tetrodotoxin to prevent action potentials. mEPSCs were detected using the template-matching algorithm in Axograph X, with a template that had 0.5 ms rise time and 5 ms decay. Statistics were computed using the Statistics Toolbox of Matlab.

### T7 Endonuclease 1 Assay and off-target analyses

Genomic PCR was performed around each sgRNA target, and related off-target sites, following the manufacturer’s instruction (using AccuPrime HiFi Taq using 500ng of genomic DNA). Products were then purified using Wizard SV Gel and PCR Clear-Up System (Promega), and quantified using a Qubit 2.0 (Thermo Fischer). T7E1 assay was performed according to manufacturer’s instructions (New England Biolabs). Briefly, 200ng of genomic PCR was combined with 2μL of NEBuffer 2 (New England Biolabs) and diluted to 19μL. Products were then hybridized by denaturing at 95 °C for 5 minutes then ramped down to 85 °C at -2°C/second. This was followed by a second decrease to 25°C at -0.1°C/second. To hybridized product, 1 μL T7E1 (M0302, New England Biolabs) was added and mixed well followed by incubation at 37°C for 15 minutes. Reaction was stopped by adding 1.5 μL of 0.25M EDTA. Products were analyzed on a 3% agarose gel and quantified using a Gel Doc XR system (BioRad). Off-target sites were identified and scored using Benchling (www.benchling.com). The top 5 off-target sites – chosen on the basis of raw score and irrespective of being in a coding region – were identified and analyzed using T7E1 assay as previously described. For TIDE (Tracking of Indels by DEcomposition) analyses (Brinkman et al., 2014), PCR was performed on genomic DNA using Accuprime Taq HiFi (Thermo Fischer) according to manufacture specifications. Briefly, reactions were cycled at 2 min at 94°C followed by 35 cycles of 98°C for 30 seconds, 58°C for 30 seconds, and 68°C for 2 minutes 30 seconds and a final extension phase of 68°C for 10 minutes. Products were then subjected to Sanger Sequencing and analyzed using the TIDE platform (https://tide.nki.nl/). The primers used for TIDE analyses are listed in **Supp. Table 1**.

### Deep Sequencing Sample Preparation and data analysis

Genomic PCR was performed using AccuPrime HiFi Taq (Life Technologies) following manufacturer’s instructions. About 200-500 ng of genomic DNA was used for each PCR reaction. Products were then purified using AMPure XP magnetic bead purification kit (Beckman Coulter) and quantified using a Nanodrop2000. Individual samples were pooled and run on an Illumina HiSeq2500 High Throughput at a run length of 2x125bp. A custom python script was developed to perform sequence analysis. For each sample, sequences with frequency of less than 100 reads were filtered from the data. Sequences in which the reads matched with primer and reverse complement subsequences classified as target sequences. These sequences were then aligned with corresponding wildtype sequence using global pairwise sequence alignment. Sequences that were misaligned through gaps or insertions around the expected cut site were classified as NHEJ events. The frequency, length, and position of matches, insertions, deletions, and mismatches were all tracked in the resulting aligned sequences.

### Statistical analysis

Statistical analysis was performed and plotted using Prism software. Student’s t-test (unpaired) or one-way ANOVA test was used to compare two or more groups respectively. A P-value <0.05 was considered significant.

### Accession Numbers

Raw reads from sequencing will be available at NCBI Bioproject PRJNA417829 upon publication.

## ACKNOWLEDGEMENTS

Methods are reported in the Supplement. Raw reads from sequencing will be available at NCBI Bioproject PRJNA417829 upon publication. We thank Sue Yeon Yi (UW-Madison) for help with constructs; and Karen Jansen-West and Lillian Daughrity (Mayo Clinic, Jacksonville) for AAV packaging and purification. This work was supported by NIH grants (R01AG048218 and R21 AG052404), and a “UW2020 grant” from the University of Wisconsin-Madison to SR.

## AUTHOR CONTRIBUTIONS

J. Sun and S. Roy designed the overall experiments, analyzed the data and assembled the manuscript; J. Sun performed most of the experiments. J. Carlson-Stevermer and K. Saha did most of the genomic analyses and analyzed the data. U. Das (UCSD) generated some constructs and performed live cell trafficking experiments in cultured neurons. M. Shen and X. Zhao (UW-Madison) planned/executed the hippocampal injection experiments; and M. Delenclos and P. McLean (Mayo Clinic, Jacksonville) planned/executed the intracerebroventricular injection experiments. A. Snead and A. Sproul (Columbia) planned and executed the iPSC experiments. L. Wang and J. Loi helped with cell culture and some data analyses. A. Petersen, M. Stockton, and A. Bhattacharyya planned/executed experiments with human embryonic stem cells; M. Jones did electrophysiology in cultured neurons. The overall idea was conceived by S. Roy.

## EXTENDED DATA (4 figures, 2 supplementary tables) Extended

**Extended Data Fig. 1.**
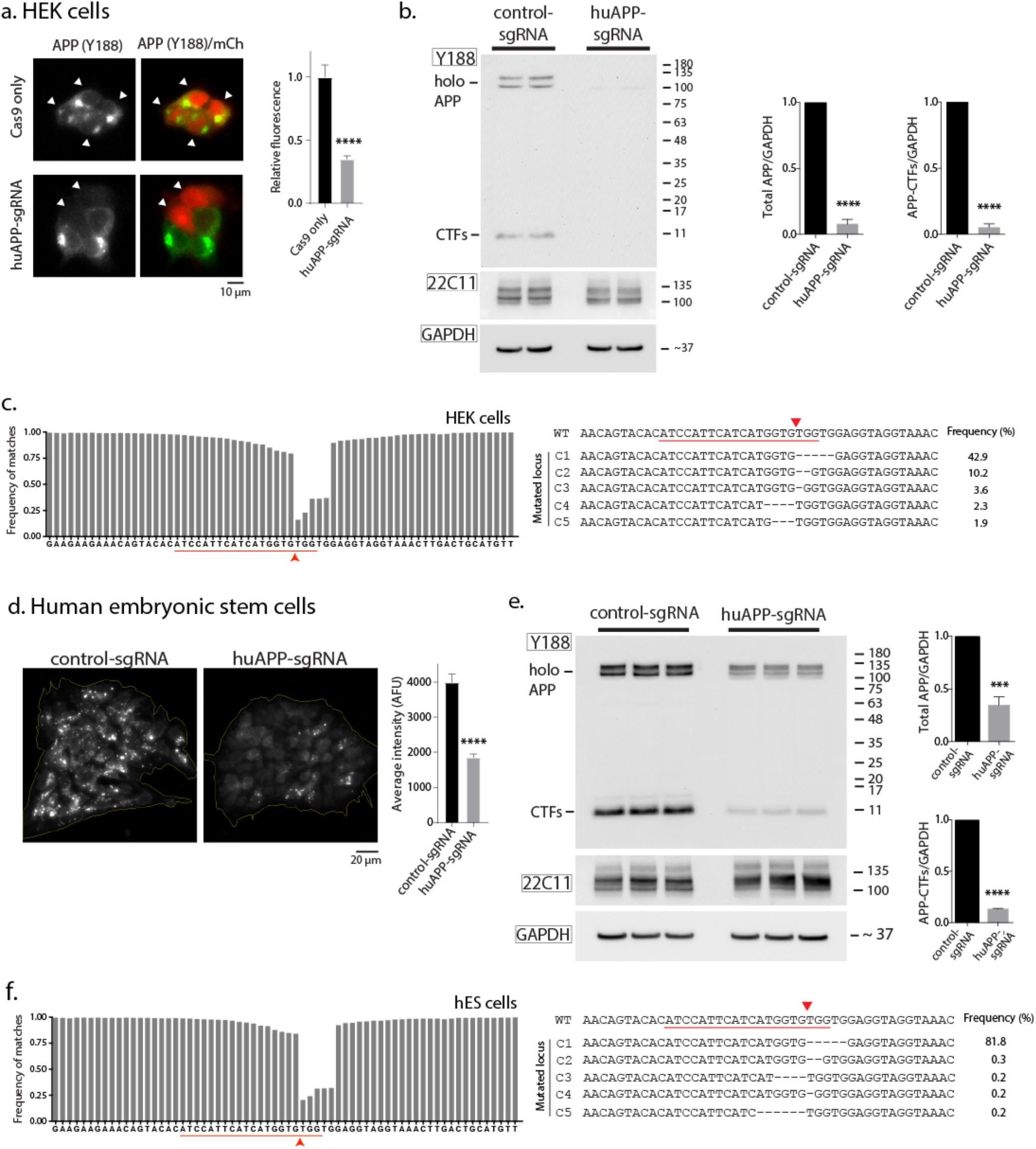
APP C-terminus editing in human cells. **(a)** HEK cells were transfected with human-specific APP-sgRNA and Cas9 (or Cas9 only), and immunostained with the Y188 antibody (after 5 days in culture). Note attenuation of staining, quantified on right (mean ± SEM of ~30 cells from two independent experiments). **(b)** HEK cells were transduced by lentiviral vectors carrying APP-sgRNA and Cas9 (or non-targeting control-sgRNA/Cas9 as control), and immunoblotted with the Y188 and 22C11 antibodies (in the presence of GSI). Note attenuation of APP-CTFs in APP-sgRNA treated cells, indicating CRISPR-editing (mean ± SEM of three independent experiments). **(c)** Deep sequencing of APP C-terminus in HEK; treated as described in (b). Red underline marks the sgRNA target sequence and arrowhead denotes predicted cut-site. Note extensive mismatch around predicted cut-site, indicating robust editing. Right: Major mutated APP loci resulting from CRISPR-editing, and their frequencies. **(d and e)** Human ESCs were transduced by lentiviral vectors carrying human APP-sgRNA/Cas9 (or non-targeting sgRNA/Cas9). Samples were immunostained with the Y188 antibody (d) or immunoblotted with the Y188 and 22C11 antibodies (e). Note attenuation of APP-CTFs in sgRNA-transduced group (mean ± SEM of ~20 colonies from two independent experiments). **(f)** Deep sequencing of APP C-terminus in human ESCs. Red underline marks the sgRNA target sequence and arrowhead denotes predicted cut-site. Note extensive mismatch around predicted cut-site, indicating robust editing. Right: Major mutated APP loci resulting from CRISPR-editing, and their frequencies.

**Extended Data Fig. 2.**
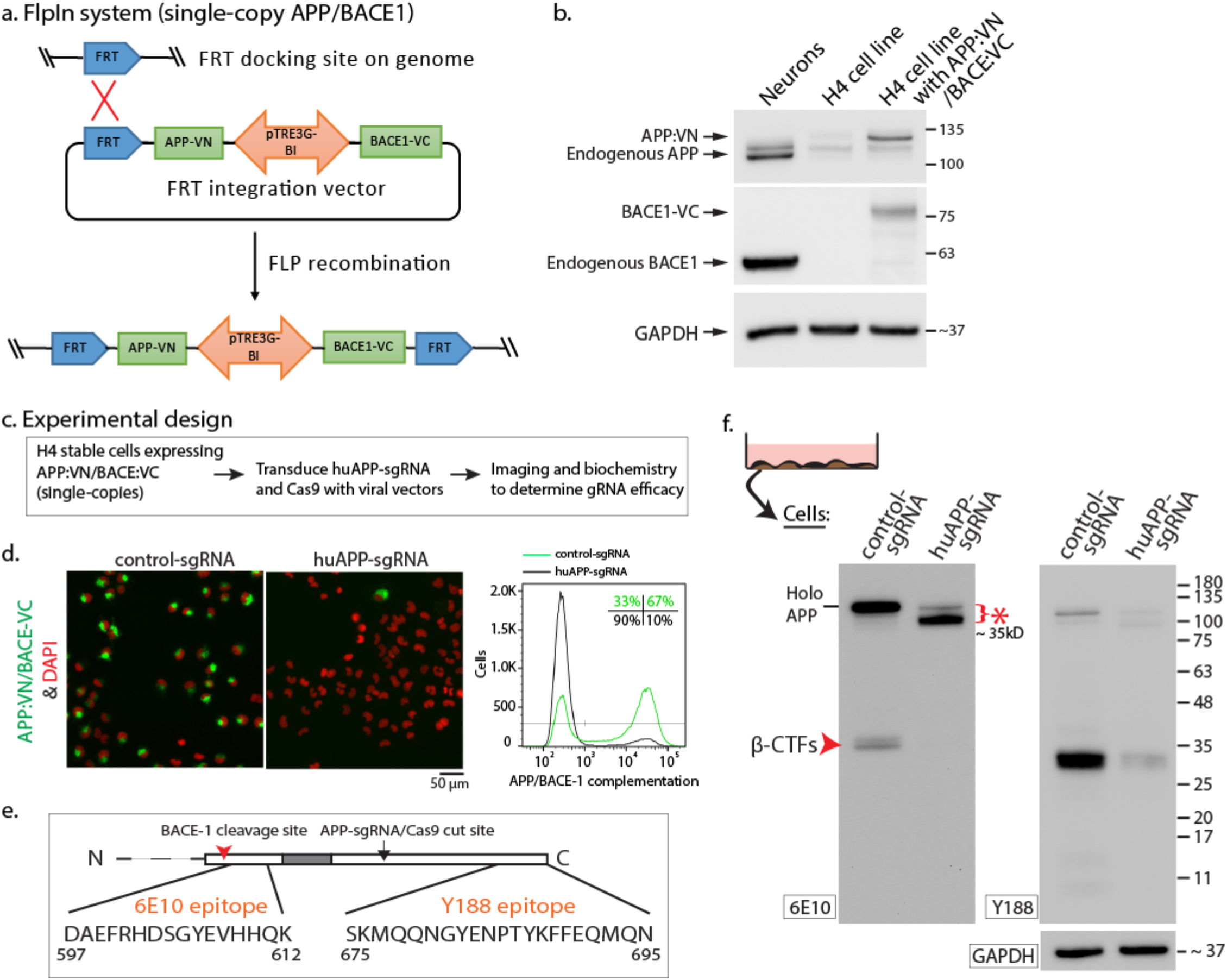
Engineering an APP/BACE-1 single copy H4 neuroglioma cell line to evaluate amyloid pathway. **(a)** Strategy to integrate APP:VN and BACE-1:VC into the H4 genome and creating a cell line expressing single copies of the two proteins (see methods for details). **(b)** APP and BACE-1 expression in the H4 cell line. Note negligible expression of endogenous proteins in native H4 cells. **(c)** Schematic of experiments. Stable H4 cell line (from a) was transduced with lentiviral vectors carrying APP-sgRNA/Cas9 (or control sgRNA/Cas9). APP/BACE-1 Venus complementation and APP cleavage products were analyzed. **(d)** The APP/BACE-1 Venus complementation visualized by fluorescence microscopy (left) or flow cytometry (right) in the H4 APP/BACE-1 VN/VC stable cell line. Note attenuation of complementation, indicating editing by the APP-sgRNA. **(e)** APP epitopes recognized by the Y188 and 6E10 antibodies. Note that in the scenario where APP is edited by CRISPR and BACE-1-cleavage is attenuated, the 6E10 antibody would only detect a fragment that is ~ 25 kD smaller than holo-APP. **(f)** Western blots of intracellular β-cleavage products from the H4 APP/BACE-1 VN/VC stable cell line above (GSI added). In the 6E10 blot, note that the fragment marked by red arrowhead in the control sample is β-CTF, which is attenuated in the APP-sgRNA treated cells.

**Extended Data Fig. 3.**
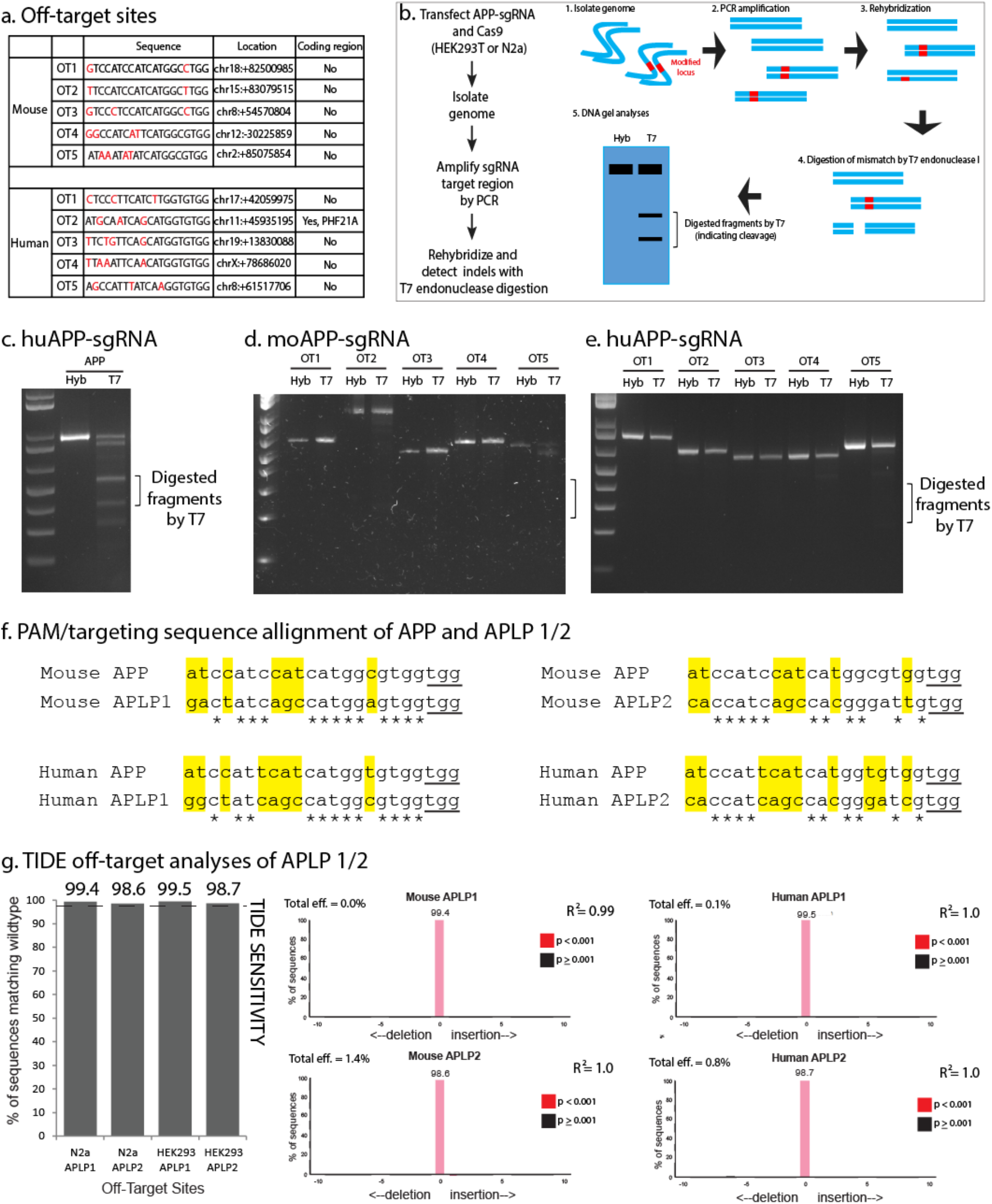
Off target analyses of APP-sgRNA. **(a)** Computationally predicted top five off-target (OT) sites in the genome, that can be potentially targeted by the mouse and human APP-sgRNAs (mismatched nucleotides in the targeting sequence are marked in red). Genomic locations corresponding to the sequences is shown on the right column (note most are in non-coding regions). **(b)** Strategy of T7 endonuclease digestion assay to detect genome-editing events. Genomic DNA was PCR amplified with primers bracketing the modified locus. PCR products were then rehybridized, yielding three possible structures. Duplexes containing a mismatch were digested by T7 endonuclease I. DNA gel analysis was used to calculate targeting efficiency. Note digested fragments in the gel indicates cleavage. **(c)** Gene edits at the APP locus by the APP-sgRNA, as seen by T7 endonuclease digestion. Note two digested fragments were recognized after T7 endonuclease digestion. **(d and e)** T7 endonuclease assays of potential off-target sites (mouse and human). No digested fragments are seen, indicating that the sgRNAs do not generate detectable gene edits at these sites. **(f)** Comparison of APLP1 and 2 sequences with APP at the sgRNA targeting site. Asterisks mark conserved nucleotide sequences, and the PAM sites are underlined. Nucleotide mismatches are highlighted in yellow. Note extensive mis-match of the mouse and human sequences at the sgRNA targeting site. **(g)** Left: Off-target TIDE analysis of APP family members APLP1 and 2 in mouse (neuro 2a) and human (HEK) cell lines following lentiviral integration of Cas9 using TIDE. No modifications were detected below the TIDE limit of detection (dotted line) in either of the populations, indicating that the APP-gRNA was unable to edit APLP 1/2. Right: TIDE analysis of APLP1 and 2 loci in mouse and human cell lines. Neither of the populations had significant editing at either of the two loci, and all sequences had a near perfect correlation to the model.

**Extended Data Fig. 4.**
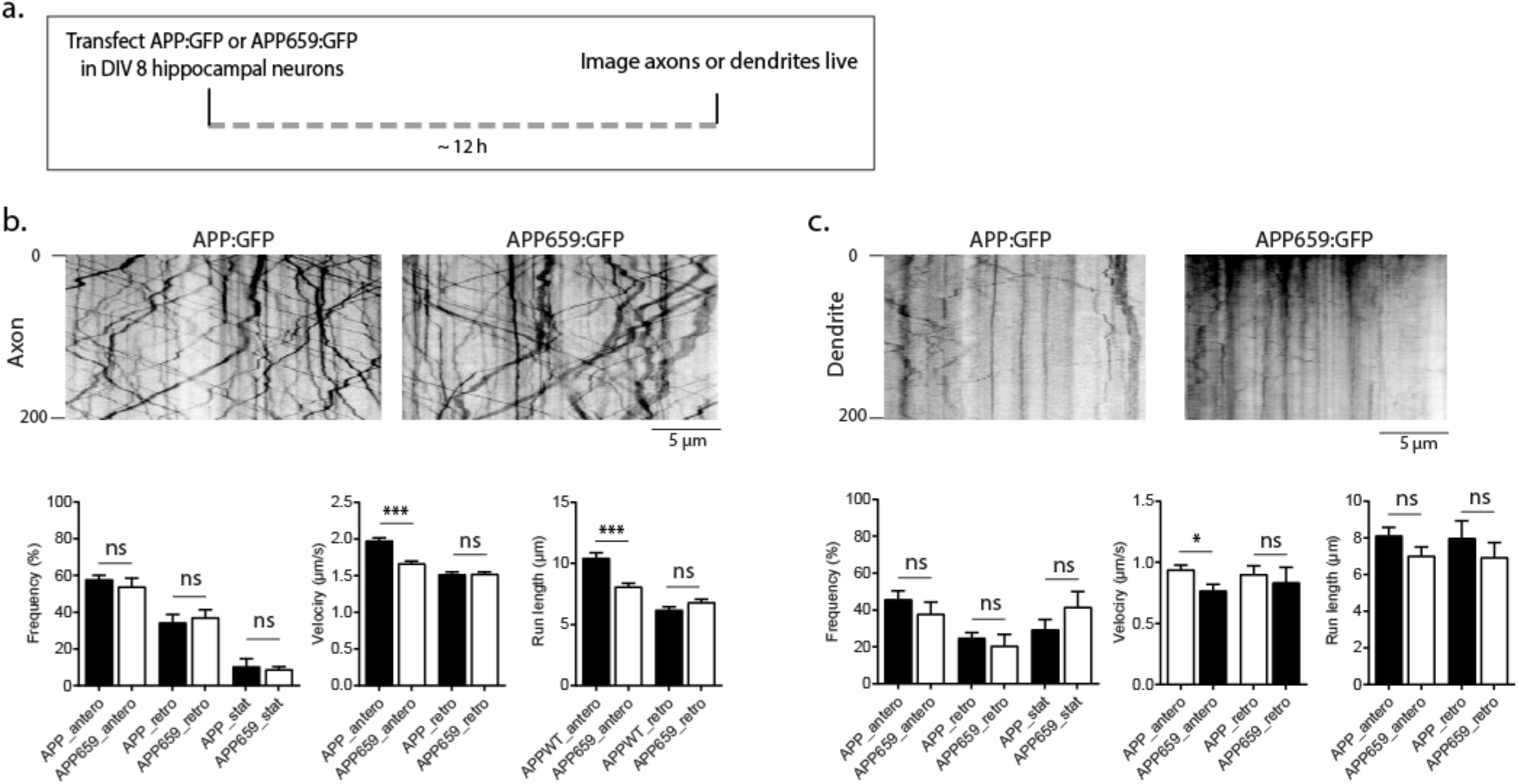
Trafficking of vesicles carrying APP(WT) or APP(659). **(a)** Cultured hippocampal neurons were transfected with APP(WT):GFP or APP(659):GFP, and kinetics of APP particles were imaged live in axons and dendrites. **(b)** Representative kymographs and quantification of APP kinetics in axons. Note that there was no change in frequency of transport, and only a modest reduction in run-length and velocity. Error bars, mean ± SEM of 325 APP(WT):GFP and 310 APP(659):GFP vesicles in 10-12 neurons from two independent experiments. **(c)** Representative kymographs and quantification of APP kinetics in dendrites. Error bars, mean ± SEM of 130 APP(WT):GFP and 115 APP(659):GFP particles in 10-12 neurons from two independent experiments.

## Sun er al., Supp. Table 1

**Table S1.**
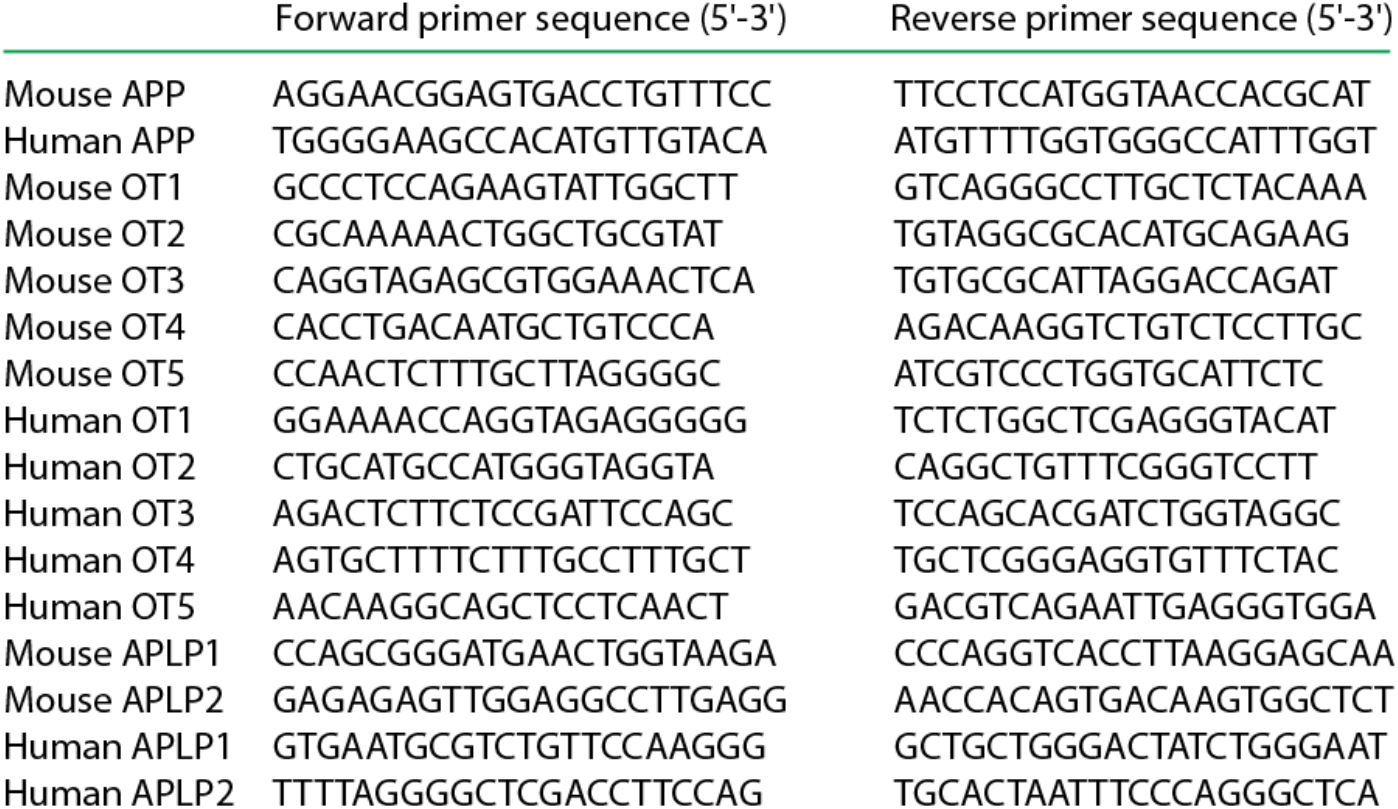
PCR primers used for on- and off-target genomic loci amlification

**Table S2.**
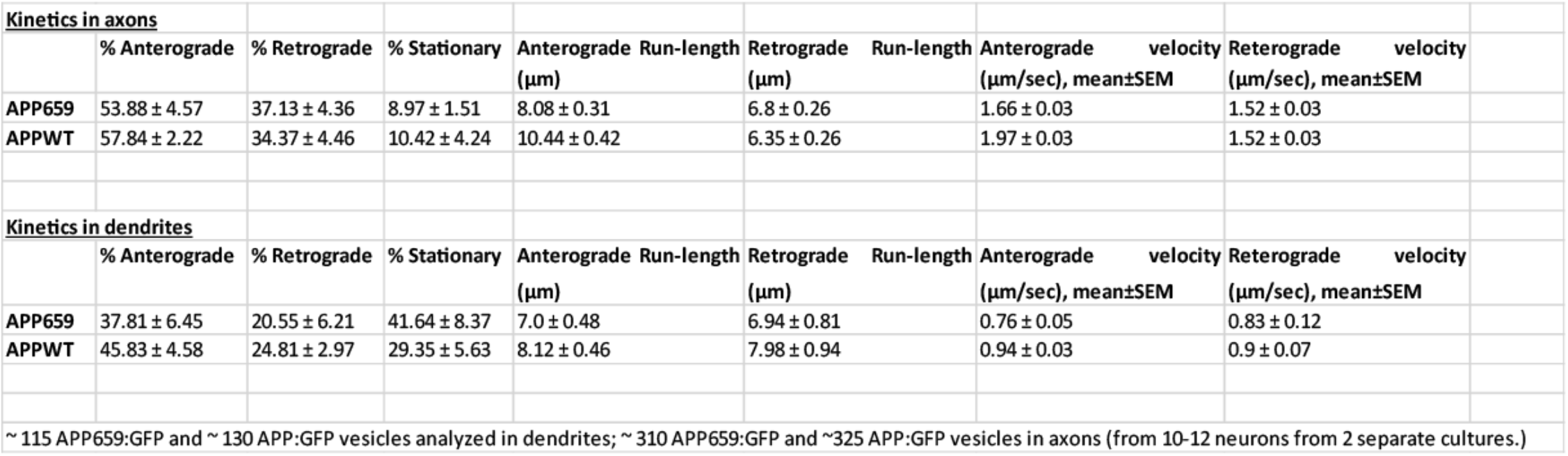
Transport parameters ofWT and 659 (“CRISPR-m¡m¡c) APP

## REFERENCES

1 Hardy, J. & Selkoe, D. J. The amyloid hypothesis of Alzheimer’s disease: progress and problems on the road to therapeutics. Science 297, 353–356 (2002).

2 De Strooper, B. & Karran, E. The Cellular Phase of Alzheimer’s Disease. Cell 164, 603–615, doi:10.1016/j.cell.2015.12.056 (2016).

3 Musiek, E. S. & Holtzman, D. M. Three dimensions of the amyloid hypothesis: time, space and ‘wingmen’. Nat Neurosci 18, 800–806, doi:10.1038/nn.4018 (2015).

4 Fellmann, C., Gowen, B. G., Lin, P. C., Doudna, J. A. & Corn, J. E. Cornerstones of CRISPR-Cas in drug discovery and therapy. Nat Rev Drug Discov 16, 89–100, doi:10.1038/nrd.2016.238 (2017).

5 Sander, J. D. & Joung, J. K. CRISPR-Cas systems for editing, regulating and targeting genomes. Nat Biotechnol 32, 347–355, doi:10.1038/nbt.2842 (2014).

6 Dunbar, C. E. et al. Gene therapy comes of age. Science 359, doi:10.1126/science.aan4672 (2018).

7 McMahon, M. A. & Cleveland, D. W. Gene therapy: Gene-editing therapy for neurological disease. Nat Rev Neurol 13, 7–9, doi:10.1038/nrneurol.2016.190 (2017).

8 Yang, S. et al. CRISPR/Cas9-mediated gene editing ameliorates neurotoxicity in mouse model of Huntington’s disease. J Clin Invest 127, 2719–2724, doi:10.1172/JCI92087 (2017).

9 Park, C. Y. et al. Reversion of FMR1 Methylation and Silencing by Editing the Triplet Repeats in Fragile X iPSC-Derived Neurons. Cell Rep 13, 234–241, doi:10.1016/j.celrep.2015.08.084 (2015).

10 Veres, A. et al. Low incidence of off-target mutations in individual CRISPR-Cas9 and TALEN targeted human stem cell clones detected by whole-genome sequencing. Cell Stem Cell 15, 27–30, doi:10.1016/j.stem.2014.04.020 (2014).

11 Doench, J. G. et al. Optimized sgRNA design to maximize activity and minimize off-target effects of CRISPR-Cas9. Nat Biotechnol 34, 184–191, doi:10.1038/nbt.3437 (2016).

12 Kuscu, C., Arslan, S., Singh, R., Thorpe, J. & Adli, M. Genome-wide analysis reveals characteristics of off-target sites bound by the Cas9 endonuclease. Nat Biotechnol 32, 677–683, doi:10.1038/nbt.2916 (2014).

13 Haass, C., Kaether, C., Thinakaran, G. & Sisodia, S. Trafficking and Proteolytic Processing of APP. Cold Spring Harb Perspect Med 2, a006270, doi:10.1101/cshperspect.a006270a006270 [pii].

14 Das, U. et al. Visualizing APP and BACE-1 approximation in neurons yields insight into the amyloidogenic pathway. Nat Neurosci 19, 55–64, doi:10.1038/nn.4188 (2016).

15 Muller, U. C. & Zheng, H. Physiological functions of APP family proteins. Cold Spring Harb Perspect Med 2, a006288, doi:10.1101/cshperspect.a006288 (2012).

16 Muller, U. C., Deller, T. & Korte, M. Not just amyloid: physiological functions of the amyloid precursor protein family. Nat Rev Neurosci 18, 281–298, doi:10.1038/nrn.2017.29 (2017).

17 Carlson-Stevermer, J. et al. Assembly of CRISPR ribonucleoproteins with biotinylated oligonucleotides via an RNA aptamer for precise gene editing. Nat Commun 8, 1711, doi:10.1038/s41467-017-01875-9 (2017).

18 Deyts, C., Thinakaran, G. & Parent, A. T. APP Receptor? To Be or Not To Be. Trends Pharmacol Sci 37, 390–411, doi:10.1016/j.tips.2016.01.005 (2016).

19 Beckett, C., Nalivaeva, N. N., Belyaev, N. D. & Turner, A. J. Nuclear signalling by membrane protein intracellular domains: the AICD enigma. Cell Signal 24, 402–409, doi:10.1016/j.cellsig.2011.10.007 (2012).

20 Pardossi-Piquard, R. & Checler, F. The physiology of the beta-amyloid precursor protein intracellular domain AICD. J Neurochem 120 Suppl 1, 109–124, doi:10.1111/j.1471-4159.2011.07475.x (2012).

21 Passini, M. A. & Wolfe, J. H. Widespread gene delivery and structure-specific patterns of expression in the brain after intraventricular injections of neonatal mice with an adeno-associated virus vector. J Virol 75, 12382–12392, doi:10.1128/JVI.75.24.12382-12392.2001 (2001).

22 Kim, J. Y., Grunke, S. D. & Jankowsky, J. L. Widespread Neuronal Transduction of the Rodent CNS via Neonatal Viral Injection. Methods Mol Biol 1382, 239–250, doi:10.1007/978-1-4939-3271-9_17 (2016).

23 Thinakaran, G. & Koo, E. H. Amyloid precursor protein trafficking, processing, and function. J Biol Chem 283, 29615–29619, doi:R800019200 [pii] 10.1074/jbc.R800019200 (2008).

24 Vassar, R. et al. Function, therapeutic potential and cell biology of BACE proteases: current status and future prospects. J Neurochem 130, 4–28, doi:10.1111/jnc.12715 (2014).

25 Das, U. et al. Activity-induced convergence of APP and BACE-1 in acidic microdomains via an endocytosis-dependent pathway. Neuron 79, 447–460, doi:S0896-6273(13)00455-8 [pii] 10.1016/j.neuron.2013.05.035.

26 Lee, M. S. et al. APP processing is regulated by cytoplasmic phosphorylation. J Cell Biol 163, 83–95, doi:10.1083/jcb.200301115 (2003).

27 Perez, R. G. et al. Mutagenesis identifies new signals for beta-amyloid precursor protein endocytosis, turnover, and the generation of secreted fragments, including Abeta42. J Biol Chem 274, 18851–18856 (1999).

28 Lai, A., Sisodia, S. S. & Trowbridge, I. S. Characterization of sorting signals in the beta-amyloid precursor protein cytoplasmic domain. J Biol Chem 270, 3565–3573 (1995).

29 Sisodia, S. S. Beta-amyloid precursor protein cleavage by a membrane-bound protease. Proc Natl Acad Sci U S A 89, 6075–6079 (1992).

30 Mendell, J. R. et al. Single-Dose Gene-Replacement Therapy for Spinal Muscular Atrophy. N Engl J Med 377, 1713–1722, doi:10.1056/NEJMoa1706198 (2017).

31 Chan, K. Y. et al. Engineered AAVs for efficient noninvasive gene delivery to the central and peripheral nervous systems. Nat Neurosci 20, 1172–1179, doi:10.1038/nn.4593 (2017).

32 Swiech, L. et al. In vivo interrogation of gene function in the mammalian brain using CRISPR-Cas9. Nat Biotechnol 33, 102–106, doi:10.1038/nbt.3055 (2015).

33 Sanjana, N. E., Shalem, O. & Zhang, F. Improved vectors and genome-wide libraries for CRISPR screening. Nat Methods 11, 783–784, doi:10.1038/nmeth.3047 (2014).

34 Joung, J. et al. Genome-scale CRISPR-Cas9 knockout and transcriptional activation screening. Nat Protoc 12, 828–863, doi:10.1038/nprot.2017.016 (2017).

35 Topol, A., Tran, N. N. & Brennand, K. J. A guide to generating and using hiPSC derived NPCs for the study of neurological diseases. J Vis Exp, e52495, doi:10.3791/52495 (2015).

36 Aime, P. et al. Trib3 Is Elevated in Parkinson’s Disease and Mediates Death in Parkinson’s Disease Models. J Neurosci 35, 10731–10749, doi:10.1523/JNEUROSCI.0614-15.2015 (2015).

37 Tang, Y. et al. Early and selective impairments in axonal transport kinetics of synaptic cargoes induced by soluble amyloid beta-protein oligomers. Traffic 13, 681–693, doi:10.1111/j.1600-0854.2012.01340.x (2012).

38 Ubelmann, F. et al. Bin1 and CD2AP polarise the endocytic generation of beta-amyloid. EMBO Rep 18, 102–122, doi:10.15252/embr.201642738 (2017).

39 Guo, W. et al. Fragile X Proteins FMRP and FXR2P Control Synaptic GluA1 Expression and Neuronal Maturation via Distinct Mechanisms. Cell Rep 11, 1651–1666, doi:10.1016/j.celrep.2015.05.013 (2015).

40 Chakrabarty, P. et al. Capsid serotype and timing of injection determines AAV transduction in the neonatal mice brain. PLoS One 8, e67680, doi:10.1371/journal.pone.0067680 (2013).

